# Allosteric signalling paths in hemoglobin: a protein dynamics network analysis

**DOI:** 10.1101/134288

**Authors:** Emanuele Monza, George Blouin, Thomas G. Spiro, Victor Guallar

## Abstract

Hemoglobin is the paradigm of cooperative protein-ligand binding. Cooperativity is the consequence of inter-subunit allosteric communication: binding at one site increases the affinity of the others. Despite half a century of studies, the mechanism behind oxygen binding in hemoglobin is not fully understood yet. In particular, it is not clear if cooperativity arises from preferential inter-subunit channels and which residues propagate the allosteric signal from one heme to the others. In this work, the heme-heme dynamical interactions have been mapped through a network-based analysis of residue conformational fluctuations, as described by molecular dynamics simulations. In particular, it was possible to suggest which inter-subunit interactions are mostly responsible of allosteric signalling and, within each pair of subunits, which protein fragments convey such signalling process.

## Introduction

Hemoglobin is a homodimer of heterodimers that binds molecular oxygen (O_2_), but also carbon monoxide (CO) and other small ligands, cooperatively. The heterodimers are made up of one α and one β subunit (Figure 1), both containing a heme group. Each of these prosthetic groups, which is bound to the protein through the proximal histidine (H87 and H92 in the α and β chain, respectively), is the responsible of uptaking one ligand molecule. Thanks to cooperative binding, hemoglobin can efficiently bind O_2_ in the lungs, where this is abundant, carry it through the circulatory system and release it in the tissue, where O_2_ partial pressure (pO_2_) is scarce (Ferrell, 2009). This unique ability makes hemoglobin a promising candidate for blood substituent design (Fronticelli et al., 2007), although its low thermostability and auto-oxidation limit the applications (Bobofchak et al., 2003). Therefore, hemoglobin needs to be designed to overcome these limitations while keeping an eye on cooperativity, which should not be compromised. Hence, hemoglobin design is a delicate multidimensional problem (not to speak about its interaction with the plethora of physiological effectors). Unfortunately, the mechanism behind hemoglobin cooperativity is far from being fully understood despite more than half a century of research (Bellelli, 2010).

**Figure 1.**
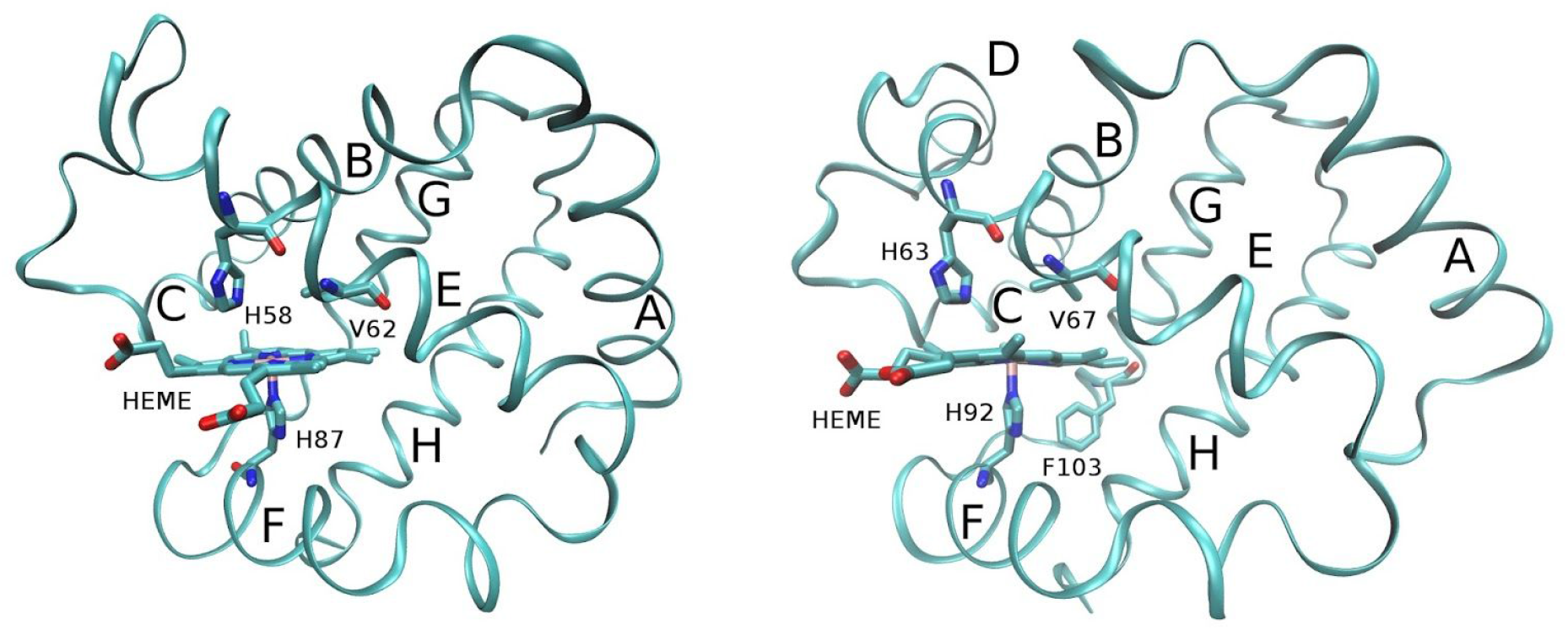
Hemoglobin’s α (left) and β (right) subunits. Helices are labeled with letters from A to H; key residues and the heme group are highlighted in licorice and labeled.

The Monod-Wyman-Changeux (MWC) model is considered one of the most reliable theories for hemoglobin cooperativity. This success was fueled by the discovery of oxy- and deoxy-hemoglobin crystal structures, which due to their different affinities were identified as the R and T states respectively (Perutz, 1970, 1979). In the deoxy-hemoglobin crystal, Fe^++^ lays out of the heme plane and the heme-His bond is tilted and elongated, while in the oxy-hemoglobin structure Fe^++^ stays in the porphyrin plane and the heme-his bond is shorter and not tilted. Both out-of-plane displacement and bond elongation are normal consequences of the high-spin (quintet) state of Fe^++^ when it is five-coordinated; however, tilting indicates that the heme-his moiety is strained. Further inspecting the crystal structures, Perutz concluded that this so called proximal strain in the deoxy-hemoglobin crystal is induced by several inter-subunit salt bridges that hold the proximal His back preventing Fe^++^ to go back to the porphyrin plane upon binding (Perutz, 1970, 1979). These salt-bridges were also predicted to break after ligand binding and trigger the T to R transition. Notably, when the heme is detached from the protein and coordinated by imidazole (analog to His side chain) (Barrick et al., 1997): (i) ligand affinity increases; (ii) the quaternary transition is inhibited and (iii) cooperativity is attenuated, but not completely lost. While the first two points validate the proximal His role in Perutz’s stereochemical model (Perutz, 1970), the last one underlines that a fraction of cooperative binding is controlled by other factors, which could comprise the configuration of the distal pocket (Olson et al., 1988) and entropic effects (Yonetani & Kanaori, 2013).

According to the tertiary two-state (TTS) model, which is a generalization of the MWC theory, ligand affinity is directly controlled by the tertiary structure of each subunit, which can exists in two states r and t with high and low affinity respectively, whose population is determined by the nature of the quaternary state (R or T) (Henry et al., 2002). In particular, almost all subunits are in the r state in R while t is predominant in T. Therefore, according to the TTS model, the quaternary state controls ligand affinity only indirectly, influencing the tertiary state of each subunit. Recently, the first computational support of the TTS model was presented by our groups (Jones et al., 2014).

The scenario turned out to be even more complicated when alternative quaternary structures were found for oxy-hemoglobin (Silva, Rogers & Arnone, 1992; Safo & Abraham, 2005). Moreover, elastic network model sampling (Xu, Tobi & Bahar, 2003) and several molecular dynamics (MD) simulations, run with different force fields (Hub, Kubitzki & de Groot, 2010; Yusuff et al., 2012; Vesper & de Groot, 2013), suggest that: (i) deoxy-hemoglobin crystal structure is very unstable in water and quickly (within 100-200 ns) converts into oxy-hemoglobin-like structures (ii) oxy-hemoglobin is well represented by a very wide ensemble of oxy-hemoglobin-like conformations instead of a few crystal structures. Nuclear magnetic resonance (NMR) (Skoog, James Holler & Crouch, 2007) experiments confirmed the second prediction (Fan et al., 2013) and showed how neither deoxy- nor oxy-hemoglobin crystal structures fit well the NMR data (Sahu et al., 2007). As a consequence, hemoglobin should be considered as a highly flexible protein described by a wide ensemble of structures, being aware that crystal structures are (only) models, with their strengths and limitations, and should not be conclusive when interpreting experimental findings. Moreover, it should be beared in mind that equilibria are dictated by free energy differences. Therefore, cooperative binding might well arise from purely dynamical effects or an altered vibrational entropy of the receptor upon ligand binding (Cooper & Dryden, 1984; Popovych et al., 2006; Hilser, Wrabl & Motlagh, 2012; Nussinov, Ruth & Chung-Jung, 2015) which could then propagate to the other protein chains.

In our previous work, it has been shown that heme reactivity is modulated by ring (out-of-plane distortion) and proximal strain (heme-His bond lengthening and tilting) in the α subunit, while only the former effect is important in the β chain (Jones et al., 2014). Such modulation can take place with no need to alter the quaternary structure of hemoglobin (Hb) and with minimal tertiary rearrangement (Jones et al., 2014). In particular, H87α, H92α (proximal and ring strain) and F103β (ring strain (Alcantara et al., 2007)) were found to have a crucial role in heme reactivity regulation. Apart from ring and proximal strain effects, affinity is also regulated in the distal pocket, where the ligand coordinates Fe (Unzai et al., 1998a). This might be more important in β since its low-affinity heme-His stretching frequency is significantly higher than the α’s despite these subunits have similar O_2_ affinities (Unzai et al., 1998b). Here, the dynamical pathways connecting these important residues (H87α, H92α, F103β, H58α, V62α, H63β and V67β) belonging to different chains are explored with molecular dynamics (MD). The aim is to understand which of these amino acids are more responsible for inter-subunit communication, how they are connected and which subunits exchange more information. In this way, the sequence search space for future stability designs (to develop better blood substituents) can be narrowed down.

## Computational Details

*System setup and molecular dynamics*. The 2dn1.pdb Hb’s R (ligated) structure was prepared with the protein preparation wizard software (Sastry et al., 2013). The protonation states of the titratable residues were assigned at pH 7 with PROPKA (Olsson et al., 2011) and double-checked with the H^++^ server (Gordon et al., 2005). After that, the following variants were prepared: (i) full-ligated (carboxy-hemoglobin, HbCO); (ii) α1 unligated; (iii) β1 unligated; full-unligated. The systems were solvated with a 10 Å buffer of waters in an orthorhombic box, neutralized and 0.15 M NaCl was added. After equilibration (default settings), 1 μs NpT production runs were performed with Desmond (Bowers et al., 2006) at 310 K for all the variants. The OPLS-2005 force-field (Kaminski et al., 2001) and the SPC explicit water model (Toukan, Kahled & Aneesur, 1985) were used. The heme parameters were generated with the hetgrp_ffgen utility of Schrödinger, inputting charges and geometries from the QM/MM calculations carried out in our previous work (Jones et al., 2014) (for the R and T states). The temperature was regulated with the Nosé−Hoover chain thermostat (Nosé & Shuichi, 1984) with a relaxation time of 1.0 ps, and the pressure was controlled with the Martyna−Tobias−Klein barostat (Martyna, Tobias & Klein, 1994) with isotropic coupling and a relaxation time of 2.0 ps. The RESPA integrator (Tuckerman, Berne & Martyna, 1992) was employed with bonded, near, and far time steps of 2.0, 2.0, and 6.0 fs, respectively. A 9 Å cutoff was used for non bonded interactions together with the smooth particle mesh Ewald method (Essmann et al., 1995).

*Allosteric signalling paths analysis*. The Weighted Implementation of Suboptimal Paths (WISP) (Van Wart et al., 2014) program was used to analyze the MD trajectories. After each residue was represented by its center of mass, the correlation (with elements C_ij_) and contact-map (with elements M_ij_) matrices were calculated. These are both NxN matrices, where N is the number of residues. Each element of the contact-map matrix is either 0 or 1 depending on the number of times the respective residues are within 4.5 Å. The functionalized correlation matrix (F) has −log|C_ij_| as elements if M_ij_ = 1 or ∞ if M_ij_ = 0. F represents a network where each residue is a node; if F_ij_ is finite, the i-th and j-th nodes are connected by an edge of distance F_ij_. The higher the absolute correlation between two residues, the lower the distance between two nodes. For any pair of nodes, a set of connecting paths can be traced, whose distance is the sum of the distances of each edge (Van Wart et al., 2014). In the seminal paper of van Wart et al., the allosteric interaction in the HisH-HisF dimer was studied; binding at the allosteric site produced a shift from 3.1-4.0 to 2.8-3.6 of the length distribution of the top 100 shortest paths that link the allosteric site to the binding pocket (Van Wart et al., 2014). Here, the top 100 shortest paths between α1-β1, α1-β2, β1-α2, α1-α2 and β1-β2 were calculated using default settings, choosing the following residues as possible end-points: F103β, H63β, V67β, H92β, H87α, H58α and V62α. While F103β, H92β and H87α account for ring and proximal strain, the remaining residues aim at describing the dynamics of the distal pocket.

## Results and Discussion

The covariance matrix of the C_α_ coordinates was built out of the superimposed oxy- (R, 2dn1) and deoxy-Hb (T, 2dn2) crystal structures and the MD trajectories were projected along its first two principal components. In such a way, the extent of R to T transition could be monitored in time. Principal component analysis (PCA) shows that, independently of the ligation state, the system is far away from reaching the T quaternary structure after 1 μs (Figure S1). Comparing the three simulations, a broader exploration by HbCO while a more pronounced R-to-T directionality can be appreciated for α1- and β1-unligated.

Despite none of the four simulations reached the T state, inter-subunit communication is differently channelled according to the WISP analysis. Common to all systems, α1-β2 (α2-β1) seems to be a privileged heme-heme signalling channel (Table 1). Then, β1-β2 signalling comes next in importance. Although intra-dimer and α1-α2 communications are comparable in α1- and β1-unligated, the former becomes more important when Hb is either saturated or lacking ligands, the latter being the least relevant allosteric channel.

**Table 1.**
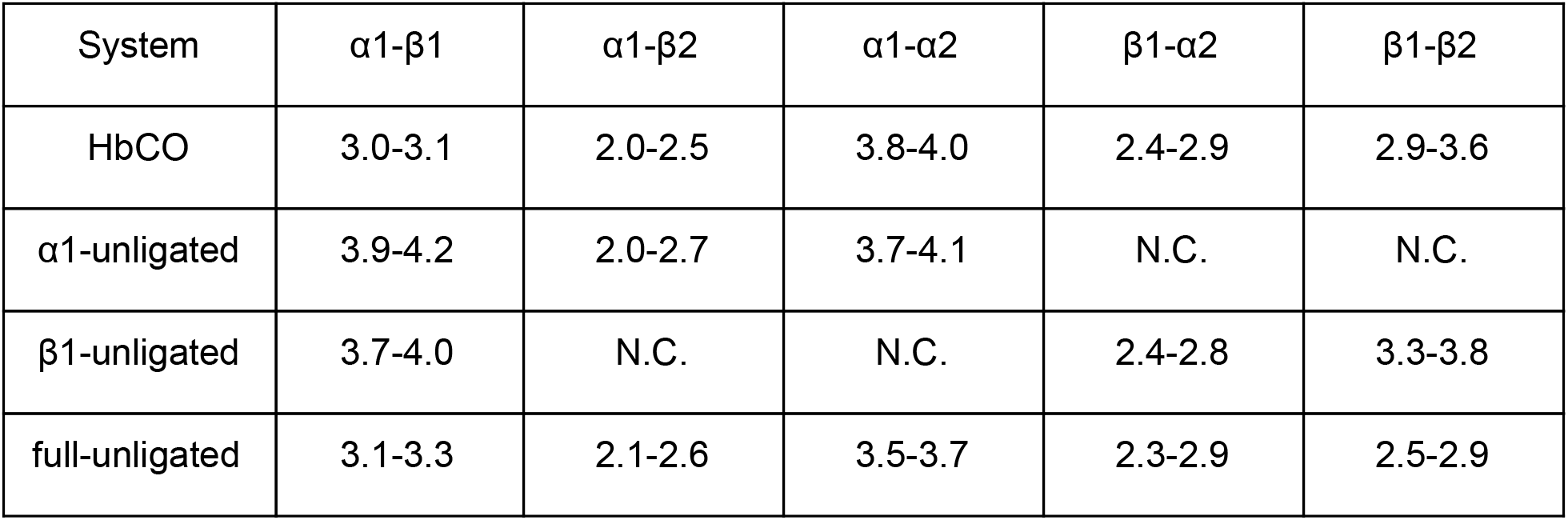
Range of lengths for the best 100 signalling paths.

### α-β Inter-dimer signalling

While α1-β2 (α2-β1) has H87α1 as sole source/sink of allosteric signal in HbCO, signalling is enriched upon CO unbinding in the α1 chain, involving also H58α1 and V62α1 (Table 2). Therefore, the response of both proximal and distal pockets to unbinding in the α1 chain are successfully propagated to the β2 subunit. When CO leaves the β1 monomer, F103β1 is the main source of the allosteric signal among the selected residues. On the other hand, β1’s distal pocket has a limited role in the β1-α2 allosteric signalling, at least in the amount of time simulated here; perhaps further quaternary rearrangement is necessary to get it more significantly involved in the β1-α2 communication. Finally, when all the subunits are unligated, the communication between proximal sites seems to dominate the scene. Regardless the ligation state, it turns out that α1-β2 (α2-β1) communication is mostly mediated by the FG corners (see Figures 8.3 and (8.4 for the illustration of the secondary structure elements) of both chains and the B, C helices of the α subunit (as a consequence of the marginality of β’s distal pocket role in heme-heme allosteric interaction, these helices are less important in this subunit), as shown in Figures 9.2-9.4. More in detail, H87α’s response to CO unbinding is transmitted to F103β through αFG-βFG correlated motion, while the dynamical response of α’s distal pocket is communicated through the αB-αC-βFG path down to F103β. When β1 loses the ligand, also the αC helix becomes crucial to convey the allosteric information to both heme’s pockets, differentiating the β-to-α and α-to-β paths.

**Figure 2.**
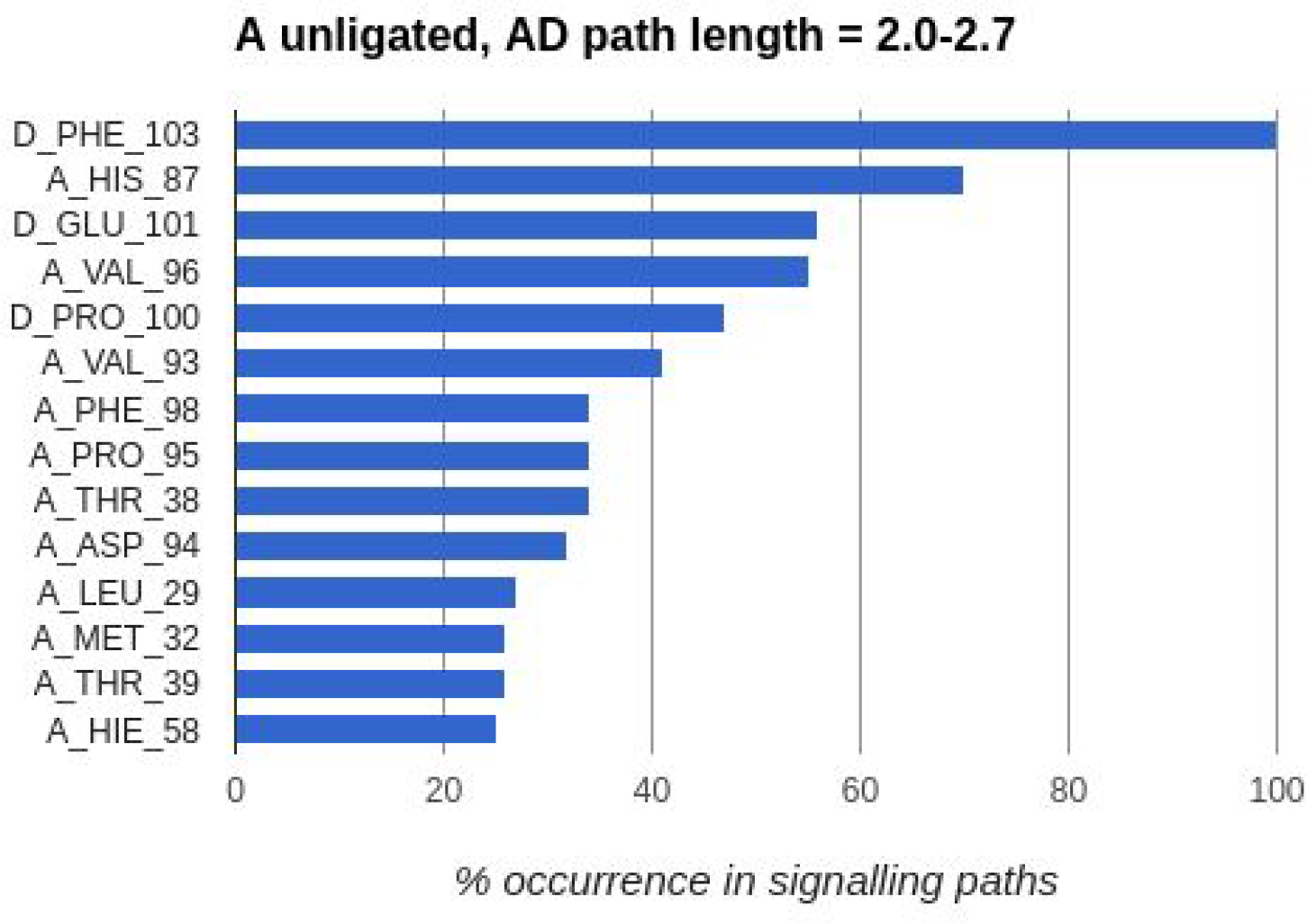
Residues that are more involved in the α1-β2 (AD) signalling in α1-uniligated. Bin heights represent the frequency at which such residues take part to the signalling process..

**Figure 3.**
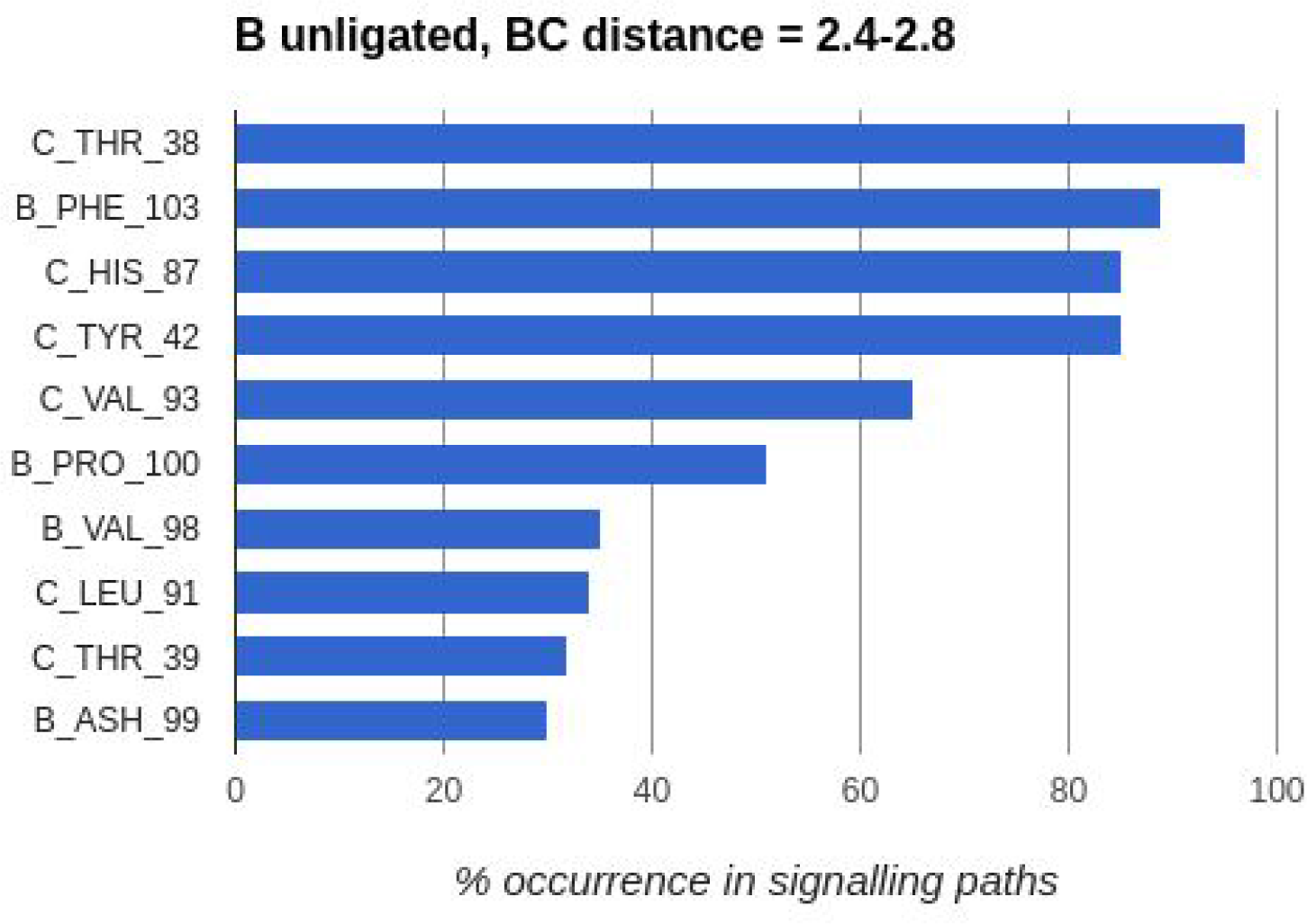
Residues that are more involved in the β1-α2 (BC) signalling in β1-uniligated. Bin heights represent the frequency at which such residues take part to the signalling process.

**Figure 4.**
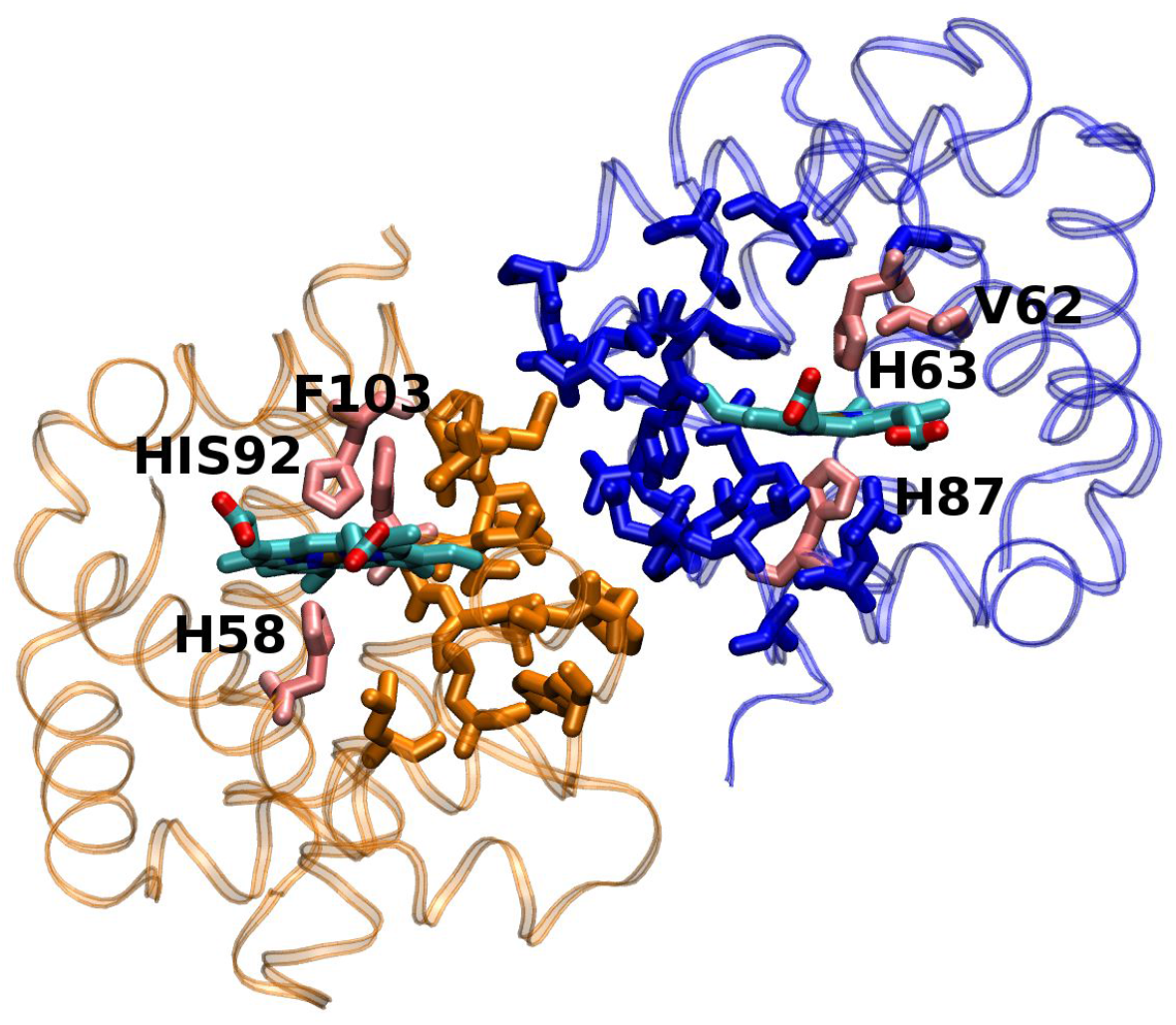
Illustration of the residues that are more involved in the α1β2 (AD) and β1α2 (BC) signalling in α1- and β1-uniligated Hb (directly obtained from Figures 9.2 and 9.3). The α chain is shown in blue, the β chain in yellow. Figure prepared with VMD (Humphrey et al., 1996) and modified with GIMP (https://www.gimp.org/).

### β-β Inter-Dimer Signalling

In such information exchange channel, F103β and H92β are the most frequent signal source/sinks (Table 2). Therefore, conversely to α1-β2 (α2-β1), H92β has a significant role in this allosteric communication route, while the distal pocket does not influence such signalling path (which seems reasonable looking at the β1-β2 relative orientation, Figure 6). The N-terminal residue seems to be crucial for β1-β2 signalling, along with the rest of the H and F helices (Figures 9.5 and 9.6).

**Figure 5.**
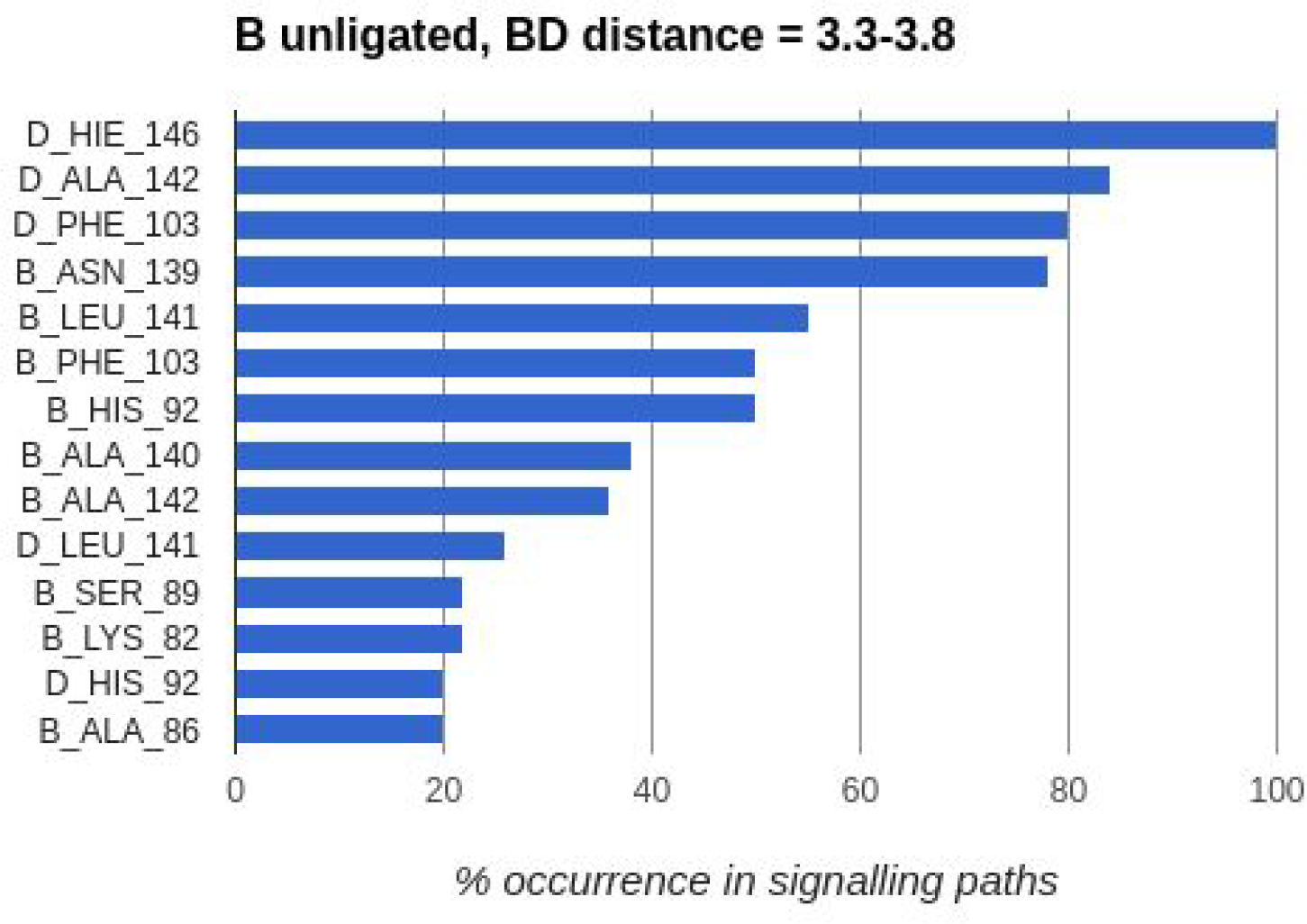
Residues that are more involved in the β1-β2 (BD) signalling in β1-unligated. Bin heights represent the frequency at which such residues take part to the signalling process.

**Figure 6.**
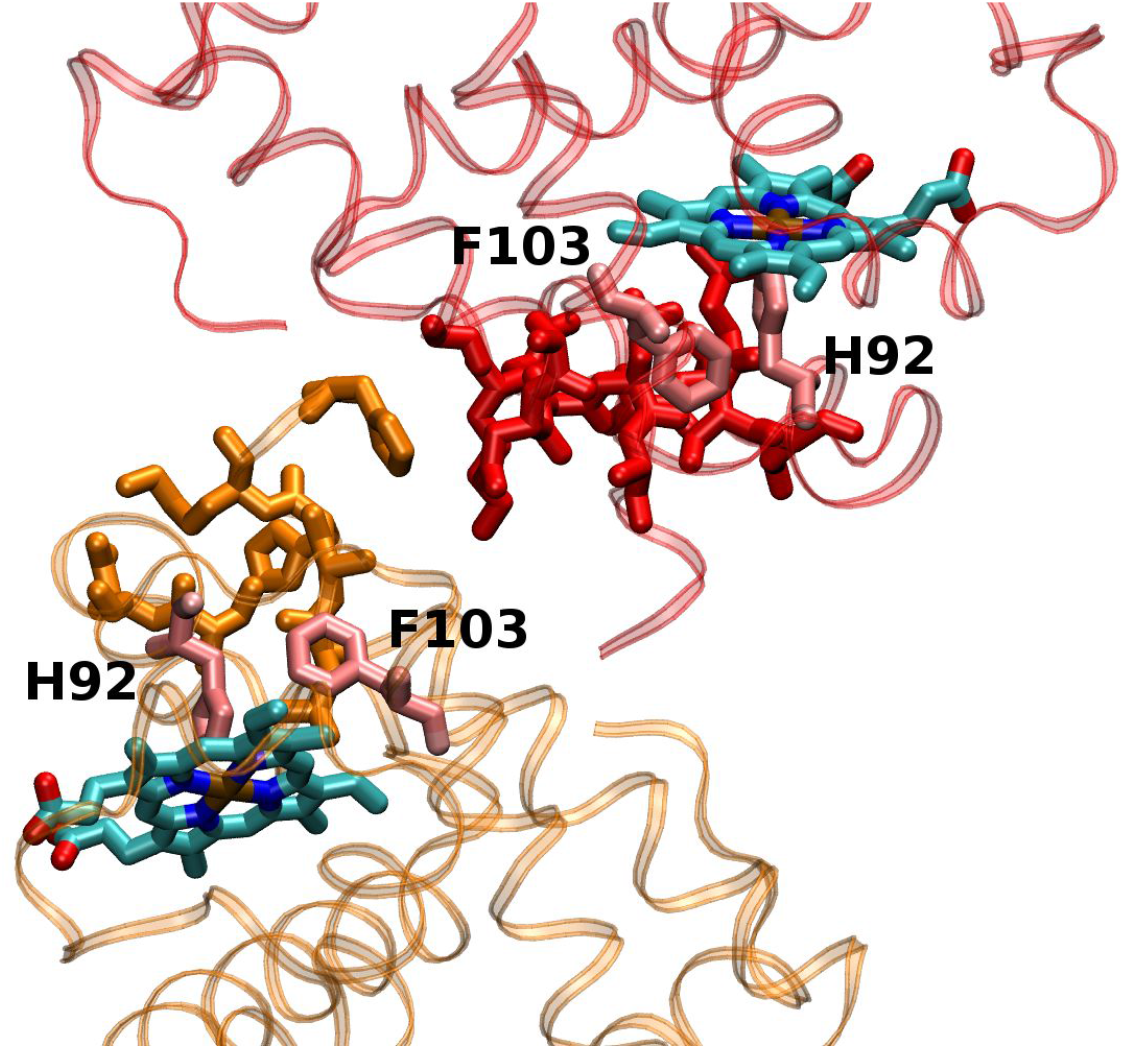
Illustration of the residues that are more involved in the β1β2 (AD) signalling in β1-unligated (directly obtained from Figure 5). The β1 chain is shown in red, the β2 chain in yellow.

### Intra-Dimer Signalling

The intra-dimer signalling paths, whose impact on cooperativity is comparable to that of the β1-β2 interaction, get more variegate upon CO unbinding from either the α1 or the β1 chain (Table 2). H87α takes an important role as signal source/sink, which is instead monopolized by the distal His when CO is bound. Therefore, Fe’s out-of-plane motion is successfully transmitted to the other chain. Moreover, H63β starts acting as a signal source/sink as well, propagating/inducing dynamical changes from/in the distal pocket.

**Figure 7.**
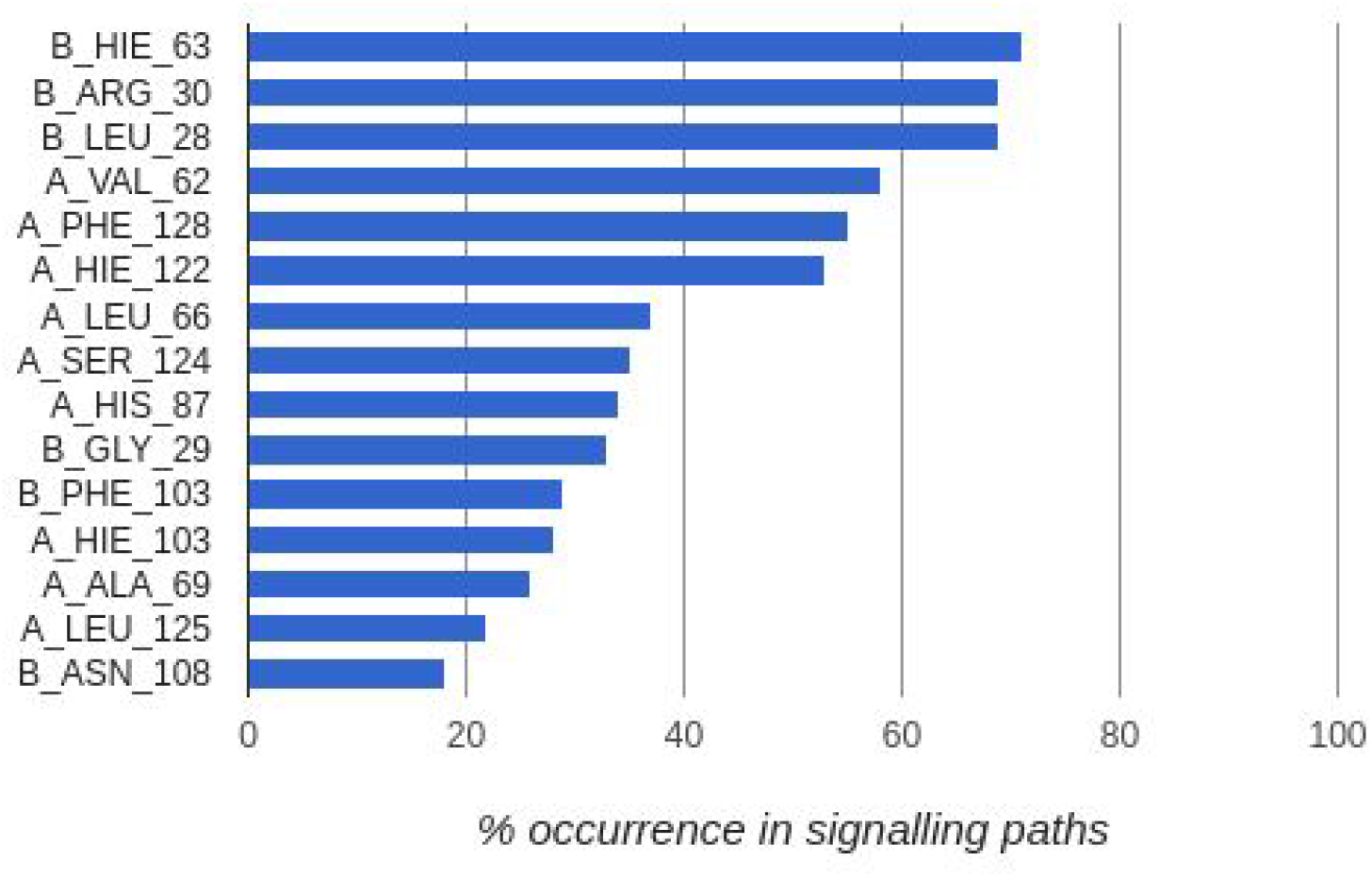
Residues that are more involved in the α1-β1 (AB) signalling in α1-unligated. Bin heights represent the frequency at which such residues take part to the signalling process.

**Figure 8.**
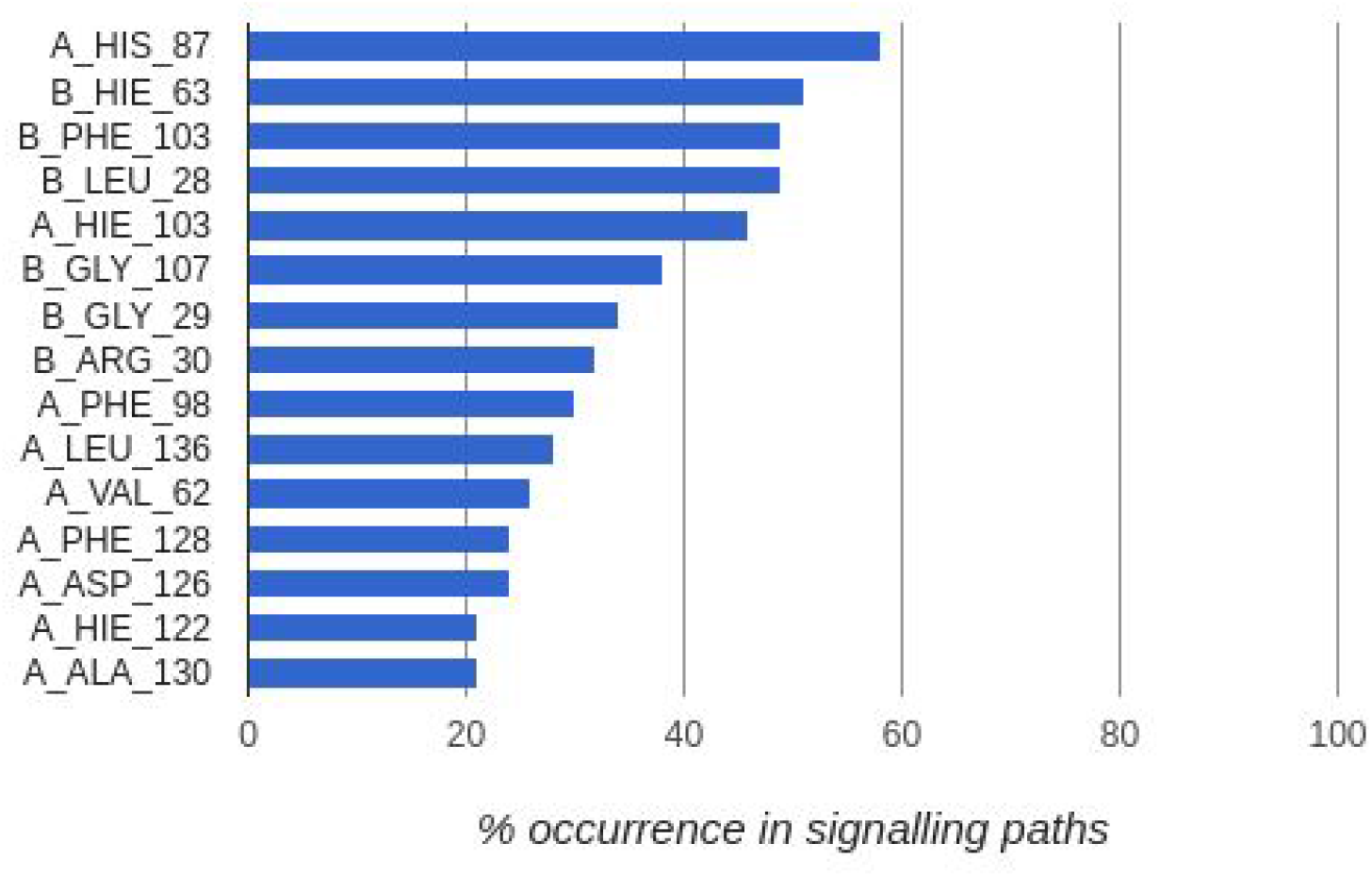
Residues that are more involved in the β1-α1 (BA) signalling in β1-unligated. Bin heights represent the frequency at which such residues take part to the signalling process.

Despite the richer number of signalling options, intra-dimer communication drops in the α1- and β1-unligated variants. Possibly, such drop might help to focus inter-subunit communication toward the most efficient allosteric channel, namely α1-β2 (α2-β1), although further studies are necessary to confirm it. However, the intra-dimer communication channel gains importance when all the subunits are unligated (Table 1), although it is still not as efficient as inter-dimer signalling is. Intra-dimer signalling is largely mediated by the following helices: αG, αE, αH, βG, βB (Figures 8 and 9). In particular, the distal pockets are mediated through the αE-αH-βB path, while H87α and F103β are linked by a αG-βG medium.

**Figure 9.**
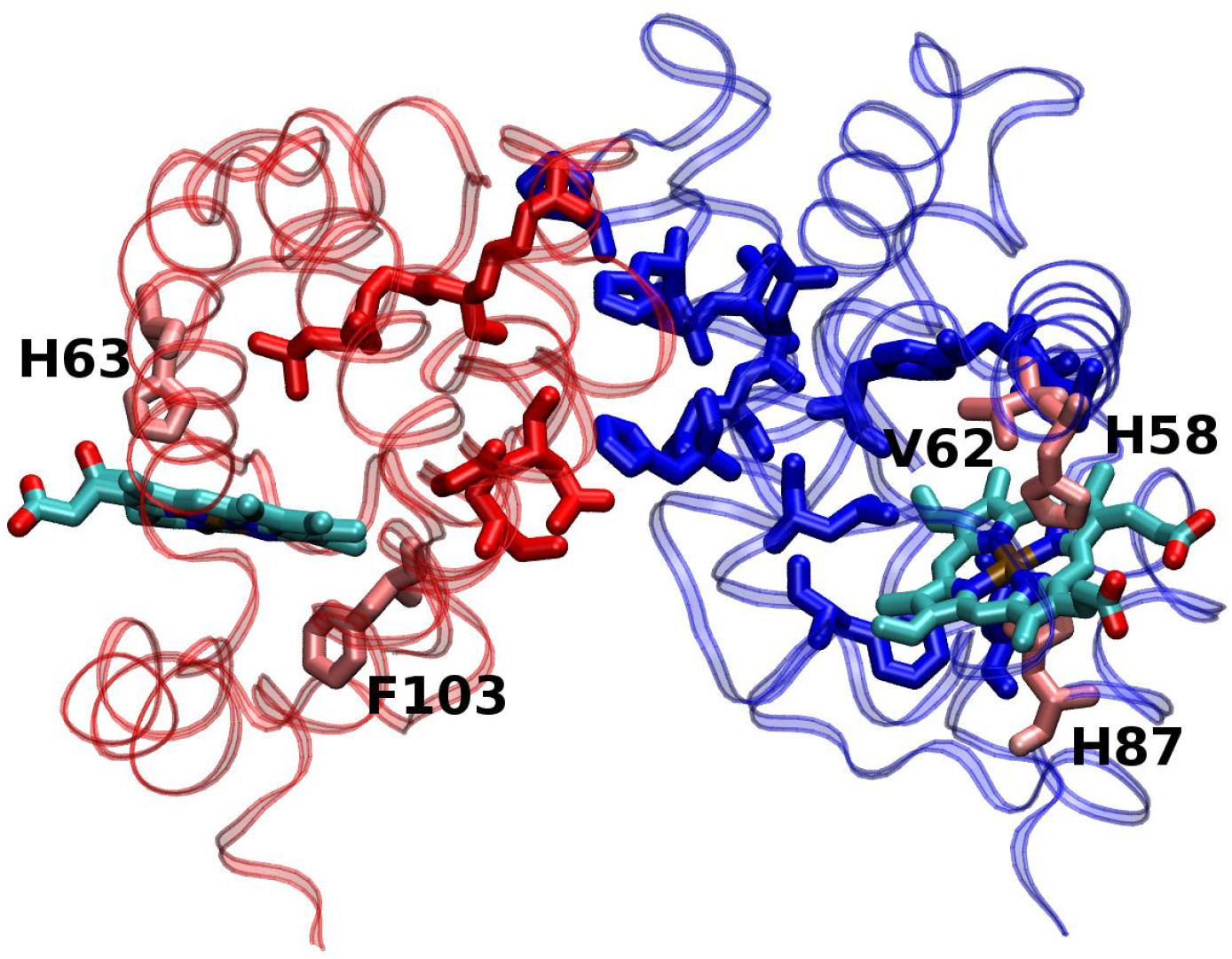
Illustration of the residues that are more involved in the α1β1 (AD) signalling in α1-uniligated (directly obtained from Figures 9.7 and 9.8). The α1 chain is shown in blue, the β1 chain in red.

**Figure 10.**
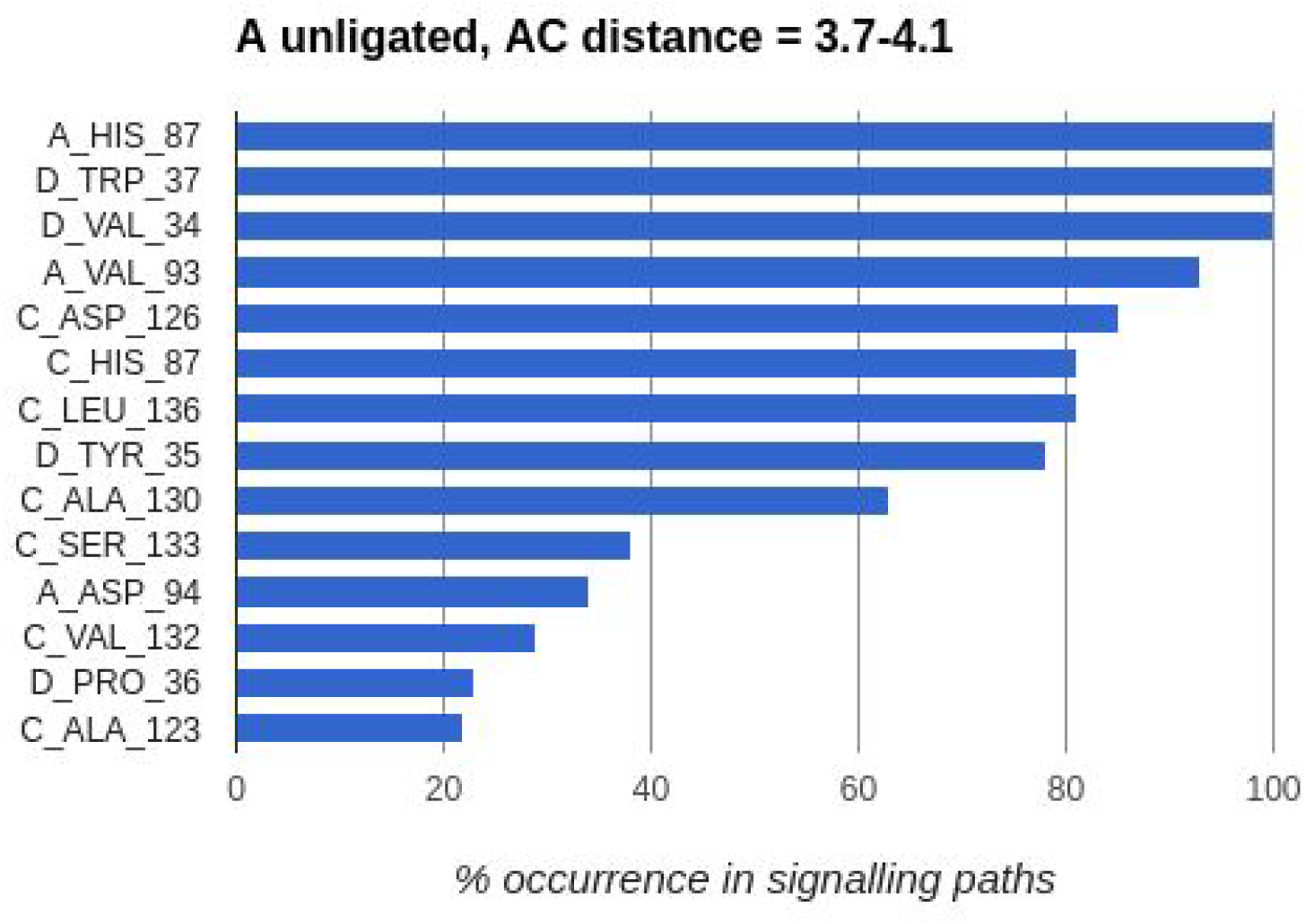
Residues that are more involved in the α1-α2 (AC) signalling in α1-unligated. Bin heights represent the frequency at which such residues take part to the signalling process.

**Figure 11.**
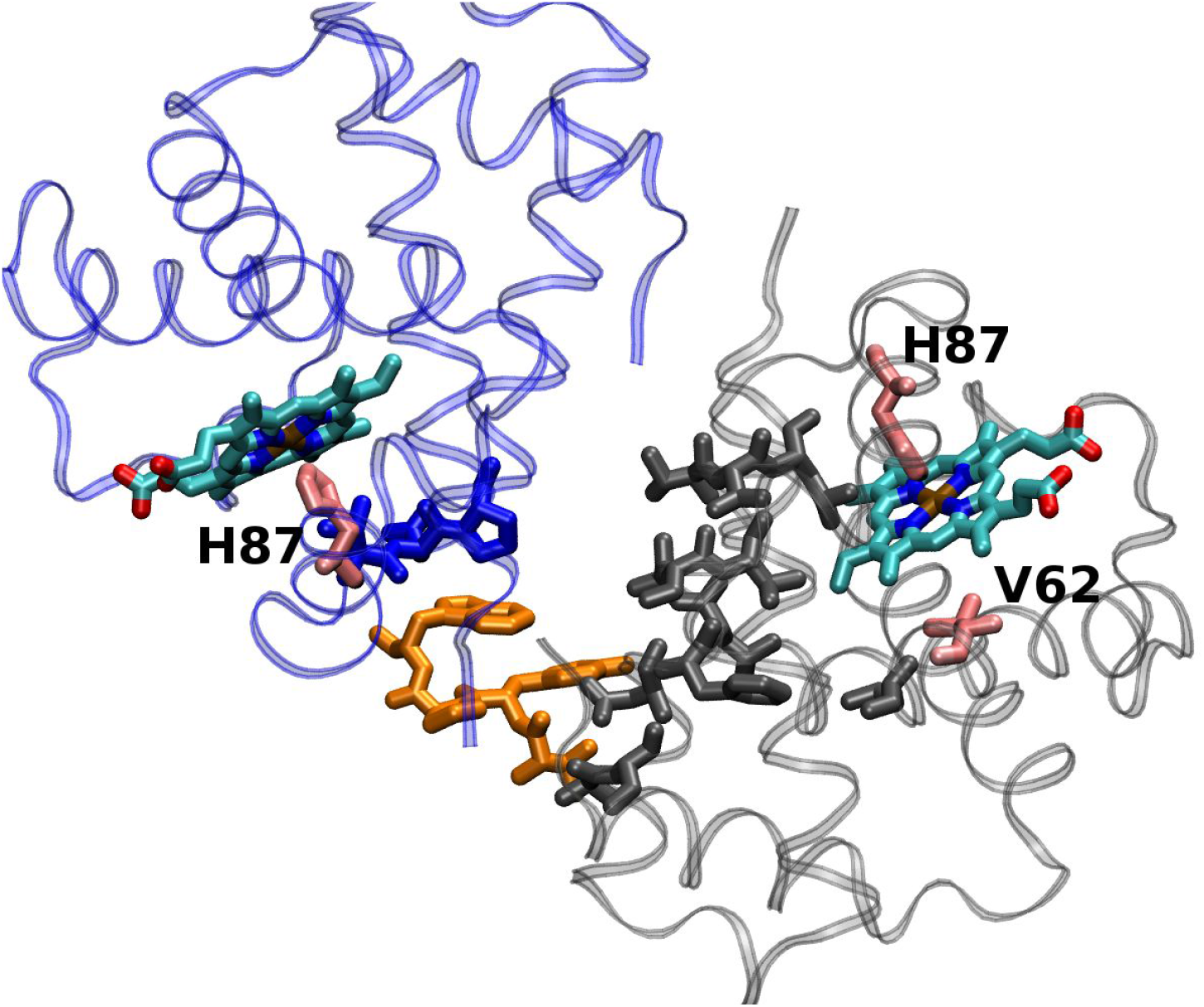
Illustration of the residues that are more involved in the α1α2 (AD) signalling in α1-uniligated (directly obtained from Figure 10). The α1 chain is shown in blue, the α2 chain in black and the β2 chain in orange. Figure prepared with VMD (Humphrey et al., 1996) and modified with GIMP (https://www.gimp.org/).

### α-α Inter-Dimer Signalling

Finally, α1-α2 communication is weak (Table 1) and mediated by a β chain (Figures 9.10, 9.11), therefore its contribution to cooperativity should be marginal. The proximal His of the ligand-free chain communicates heme’s response to the proximal and distal pockets of the other subunit through the α1FG-β2C-α2H and α1FG-β2C-α2H-α2E routes respectively. The latter, and therefore the involvement of the distal pocket, is less probable though.

## Conclusions

According to the results obtained, whether or not the first ligand unbinding event occurs in an α or β chain, the preferential route for allosteric signalling should be the inter-dimer α-β correlation path. This involves both the proximal and distal pockets of the α chain, while the β heme mostly uses F103 to communicate with the rest of the protein. In this signalling route, the following protein fragments act as bridges: αFG-βFG and αB-αC-βFG.

The β-β and intra-dimer α-β paths seem to be less important. The former connects F103 and H92 of both chains to each other through the F and H helices, the latter serves as main communication channel for β’s distal pocket. In this last case, these are the bridging moieties: αE-αH-βB and αG-βG. It should be noted that intra-dimer α-β signalling is more important in fully ligated and unligated forms than in the intermediate ligation states. Finally, α-α communication is not expected to influence cooperativity.

Collecting all the results, it can be seen that, for both chains, the following secondary structure elements are never involved in inter-subunit communication paths: the A helix and the first residues of the B helix, the EF loops and the residues within helices C and E. Therefore, their modifications for design purposes (stability in water) should not impact Hb’s cooperativity.

## REFERENCES

Alcantara RE., Xu C., Spiro TG., Guallar V. 2007. A quantum-chemical picture of hemoglobin affinity. Proceedings of the National Academy of Sciences of the United States of America 104:18451–18455.

Barrick D., Ho NT., Simplaceanu V., Dahlquist FW., Ho C. 1997. A test of the role of the proximal histidines in the Perutz model for cooperativity in haemoglobin. Nature structural biology 4:78–83.

Bellelli A. 2010. Hemoglobin and cooperativity: Experiments and theories. Current protein & peptide science 11:2–36.

Bobofchak KM., Toshiaki M., Texel SJ., Andrea B., Masaaki N., Traystman RJ., Koehler RC., Brinigar WS., Clara F. 2003. A recombinant polymeric hemoglobin with conformational, functional, and physiological characteristics of an in vivo O2transporter. American Journal of Physiology - Heart and Circulatory Physiology 285:H549–H561.

Bowers K., Kevin B., Edmond C., Huafeng X., Ron D., Michael E., Brent G., John K., Istvan K., Mark M., Federico S., John S., Yibing S., David S. 2006. Scalable Algorithms for Molecular Dynamics Simulations on Commodity Clusters. In: ACM/IEEE SC 2006 Conference (SC’06). DOI: 10.1109/sc.2006.54.

Cooper A., Dryden DT. 1984. Allostery without conformational change. A plausible model. European biophysics journal: EBJ 11:103–109.

Essmann U., Ulrich E., Lalith P., Berkowitz ML., Tom D., Hsing L., Pedersen LG. 1995. A smooth particle mesh Ewald method. The Journal of chemical physics 103:8577.

Fan J-S., Jing-Song F., Yu Z., Wing-Yiu C., Virgil S., Ho NT., Chien H., Daiwen Y. 2013. Solution Structure and Dynamics of Human Hemoglobin in the Carbonmonoxy Form. Biochemistry 52:5809–5820.

Ferrell JE Jr. 2009. Q&A: Cooperativity. Journal of biology 8:53.

Fronticelli C., Clara F., Koehler RC., Brinigar WS. 2007. Recombinant Hemoglobins as Artificial Oxygen Carriers. Artificial cells, blood substitutes, and immobilization biotechnology 35:45–52.

Gordon JC., Myers JB., Folta T., Shoja V., Heath LS., Onufriev A. 2005. H++: a server for estimating pKas and adding missing hydrogens to macromolecules. Nucleic acids research 33:W368–71.

Henry ER., Stefano B., James H., Eaton WA. 2002. A tertiary two-state allosteric model for hemoglobin. Biophysical chemistry 98:149–164.

Hilser VJ., Wrabl JO., Motlagh HN. 2012. Structural and energetic basis of allostery. Annual review of biophysics 41:585–609.

Hub JS., Kubitzki MB., de Groot BL. 2010. Spontaneous quaternary and tertiary T-R transitions of human hemoglobin in molecular dynamics simulation. PLoS computational biology 6:e1000774.

Humphrey W., William H., Andrew D., Klaus S. 1996. VMD: Visual molecular dynamics. Journal of molecular graphics 14:33–38.

Jones EM., Monza E., Balakrishnan G., Blouin GC., Mak PJ., Zhu Q., Kincaid JR., Guallar V., Spiro TG. 2014. Differential control of heme reactivity in alpha and beta subunits of hemoglobin: a combined Raman spectroscopic and computational study. Journal of the American Chemical Society 136:10325–10339.

Kaminski GA., Friesner RA., Julian T-R., Jorgensen WL. 2001. Evaluation and Reparametrization of the OPLS-AA Force Field for Proteins via Comparison with Accurate Quantum Chemical Calculations on Peptides †. The journal of physical chemistry. B 105:6474–6487.

Martyna GJ., Tobias DJ., Klein ML. 1994. Constant pressure molecular dynamics algorithms. The Journal of chemical physics 101:4177.

Nosé S., Shuichi N. 1984. A unified formulation of the constant temperature molecular dynamics methods. The Journal of chemical physics 81:511.

Nussinov R., Ruth N., Chung-Jung T. 2015. Allostery without a conformational change? Revisiting the paradigm. Current opinion in structural biology 30:17–24.

Olson JS., Mathews AJ., Rohlfs RJ., Springer BA., Egeberg KD., Sligar SG., Tame J., Renaud JP., Nagai K. 1988. The role of the distal histidine in myoglobin and haemoglobin. Nature 336:265–266.

Olsson MHM., Søndergaard CR., Rostkowski M., Jensen JH. 2011. PROPKA3: Consistent Treatment of Internal and Surface Residues in Empirical pKa Predictions. Journal of chemical theory and computation 7:525–537.

Perutz MF. 1970. Stereochemistry of cooperative effects in haemoglobin. Nature 228:726–739.

Perutz MF. 1979. Regulation of Oxygen Affinity of Hemoglobin: Influence of Structure of the Globin on the Heme Iron. Annual review of biochemistry 48:327–386.

Popovych N., Nataliya P., Shangjin S., Ebright RH., Kalodimos CG. 2006. Dynamically driven protein allostery. Nature structural & molecular biology 13:831–838.

Safo MK., Abraham DJ. 2005. The Enigma of the Liganded Hemoglobin End State: A Novel Quaternary Structure of Human Carbonmonoxy Hemoglobin †, ‡. Biochemistry 44:8347–8359.

Sahu SC., Simplaceanu V., Gong Q., Ho NT., Tian F., Prestegard JH., Ho C. 2007. Insights into the solution structure of human deoxyhemoglobin in the absence and presence of an allosteric effector. Biochemistry 46:9973–9980.

Sastry GM., Adzhigirey M., Day T., Annabhimoju R., Sherman W. 2013. Protein and ligand preparation: parameters, protocols, and influence on virtual screening enrichments. Journal of computer-aided molecular design 27:221–234.

Silva MM., Rogers PH., Arnone A. 1992. A third quaternary structure of human hemoglobin A at 1.7-A resolution. The Journal of biological chemistry 267:17248–17256.

Skoog DA., James Holler F., Crouch SR. 2007. Principles of Instrumental Analysis. Brooks/Cole Publishing Company.

Toukan K., Kahled T., Aneesur R. 1985. Molecular-dynamics study of atomic motions in water. Physical Review B: Condensed Matter and Materials Physics 31:2643–2648.

Tuckerman M., Berne BJ., Martyna GJ. 1992. Reversible multiple time scale molecular dynamics. The Journal of chemical physics 97:1990.

Unzai S., Eich R., Shibayama N., Olson JS., Morimoto H. 1998a. Rate Constants for O2 and CO Binding to the and Subunits within the R and T States of Human Hemoglobin. The Journal of biological chemistry 273:23150–23159.

Unzai S., Eich R., Shibayama N., Olson JS., Morimoto H. 1998b. Rate Constants for O2 and CO Binding to the and Subunits within the R and T States of Human Hemoglobin. The Journal of biological chemistry 273:23150–23159.

Van Wart AT., Jacob D., Lane V., Amaro RE. 2014. Weighted Implementation of Suboptimal Paths (WISP): An Optimized Algorithm and Tool for Dynamical Network Analysis. Journal of chemical theory and computation 10:511–517.

Vesper MD., de Groot BL. 2013. Collective dynamics underlying allosteric transitions in hemoglobin. PLoS computational biology 9:e1003232.

Xu C., Tobi D., Bahar I. 2003. Allosteric changes in protein structure computed by a simple mechanical model: hemoglobin TR2 transition. Journal of molecular biology 333:153–168.

Yonetani T., Kanaori K. 2013. How does hemoglobin generate such diverse functionality of physiological relevance? Biochimica et biophysica acta 1834:1873–1884.

Yusuff OK., Babalola JO., Bussi G., Raugei S. 2012. Role of the subunit interactions in the conformational transitions in adult human hemoglobin: an explicit solvent molecular dynamics study. The journal of physical chemistry. B 116:11004–11009.

